# Non-catalytic and catalytic TREHALOSE-6-PHOSPHATE SYNTHASES interact with RAMOSA3 to control maize development

**DOI:** 10.1101/2025.08.09.669499

**Authors:** Thu M Tran, Hannes Claeys, María Jazmín Abraham-Juárez, Son L Vi, Xiaosa Xu, Kevin Michalski, Tsung Han Chou, Sessen D Iohannes, Panagiotis Boumpas, Z’Dhanne P Williams, Samatha Sheppard, Cara Griffiths, Matthew J Paul, Hiro Furukawa, David Jackson

## Abstract

Trehalose-6-phosphate (Tre6P) is the intermediate in the two-step pathway of trehalose biosynthesis mediated by Tre6P-synthases (TPSs) and Tre6P-phosphatases (TPPs). Plants harbor small families of *TPS* and *TPP* genes, however most plant TPSs lack enzymatic activity, suggesting they have regulatory functions. The classical mutant *ramosa3* (*ra3*) increases inflorescence branching in maize, and RA3 encodes a catalytic TPP. We found that RA3 interacts with maize ZmTPS1, a non-catalytic TPS. Mutants in Zm*TPS1* and its close paralog Zm*TPS12* enhance *ra3* phenotypes, suggesting their physical interaction is biologically significant. ZmTPS1 also interacts with the two catalytically active maize TPSs, ZmTPS11 and ZmTPS14, however *zmtps11;zmtps14* double mutants fail to complete embryogenesis, suggesting that they are essential, as in arabidopsis. Interestingly, the non-catalytic ZmTPS1 protein stimulated the coupled activity of RA3 and ZmTPS14, suggesting that RA3, ZmTPS1, and ZmTPS14 form a complex, and we confirmed this by expressing and purifying the three proteins and by Alphafold predictions. Our results suggest that non-catalytic TPSs form a complex with catalytic TPSs and TPPs to stimulate catalytic activity and regulate plant development.

## Introduction

Trehalose was first discovered in plants in the 1960s, but research into its molecular signaling pathways only gained momentum in the early 2000s (Elbein, 2003). Initially, trehalose was considered a storage sugar and an osmotic regulator (Crowe et al., 1992). However, we now understand that trehalose-6-phosphate (Tre6P), the precursor to trehalose, is a key regulator of plant growth and development (Dijken et al., 2004; Eastmond et al., 2002a; Fichtner and Lunn, 2021; Figueroa and Lunn, 2016; Göbel and Fichtner, 2023; Schluepmann et al., 2003). Tre6P acts as a crucial sugar signal, correlating with sucrose levels and regulating various developmental processes, such as flowering and germination (Göbel and Fichtner, 2023). Tre6P also connects sugar metabolism to hormonal pathways, particularly strigolactone signaling, to control bud dormancy. For example, reducing Tre6P levels in buds by manipulating TPP expression delays bud outgrowth, highlighting its importance in shoot branching (Wang et al., 2023). Furthermore, mutations in the enzymes that mediate Tre6P metabolism, namely trehalose-6-phosphate synthases (TPSs) and trehalose-6-phosphate phosphatases (TPPs), affect specific developmental processes, such as embryonic and vegetative development, seed dormancy (Ponnu et al., 2020; Fichtner et al., 2021), flowering (Schluepmann et al., 2003; Wahl et al., 2013), meristem determinacy (Claeys et al., 2019; Fichtner et al., 2021), and cell fate specification (Han et al., 2020).

Maize (*Zea mays*) and other flowering plants encode multiple TPSs, which are categorized into class I (catalytic) or class II (non-catalytic). The class I TPS in arabidopsis, AtTPS1, is important for development, as *attps1* mutants arrest at the embryonic torpedo stage (Eastmond et al., 2002b). While many studies have focused on Class I TPS proteins, research on the non-catalytic Class II TPS proteins remains limited, leaving their roles in plant development and physiology less understood. Loss-of-function and overexpression studies of selected class II TPS genes in arabidopsis reveal stress- and development-related phenotypes. For example, *attps5* mutants are more sensitive to abscisic acid (ABA), have reduced thermotolerance, and increased susceptibility to bacterial pathogens, indicating its role in hormone signaling, heat response, and basal immunity (Suzuki et al., 2008; Tian et al., 2019; Wang et al., 2019). In wheat (Triticum aestivum L.) the class II TPS *TaTPS11* also functions in response to both abiotic and biotic stresses, since its overexpression in arabidopsis improves cold tolerance, whereas *attps11* loss-of-function mutants are more susceptible to aphid attack, indicating its roles in cold adaptation and defense against insects (Liu et al., 2019; Singh et al., 2011). *AtTPS6* is implicated in control of cell morphology, as mutants have distorted epidermal cell shapes, suggesting a role in cytoskeletal or cell wall regulation (Chary et al., 2008). These findings suggest that class II TPS proteins perform important roles in plant stress physiology and development. How these non-catalytic TPSs function in the context of Tre6P biosynthesis is largely unknown. However, they repress the activity of Sucrose Non-Fermenting 1-related protein Kinase 1 (SnRK1), a master regulator of energy homeostasis that balances metabolism and growth (Avidan et al., 2024; Van Leene et al., 2022), suggesting the class II TPSs link Tre6P signaling to energy-sensing pathways.

*RAMOSA3* (*RA3*) encodes a maize TPP enzyme that controls inflorescence architecture by regulating meristem determinacy and branching. Maize inflorescence architecture is defined by a series of branching events, and *ra3* mutants increase both tassel and ear branching. Intriguingly, a catalytically inactive RA3 variant can partially complement the mutant phenotype (Claeys et al., 2019), suggesting that RA3 and other TPPs may have “moonlighting” regulatory roles beyond their enzymatic functions. This hypothesis is supported by the localization of RA3 protein in nuclear puncta (Demesa-Arevalo et al., 2021). While much is known about the individual roles of TPSs and TPPs, the nature of their interactions is less understood. In fungi, TPS and TPP enzymes form complexes that coordinate their functions. For example, TPSs and TPP proteins interact in *Candida albicans*, and mutations in these components disrupt trehalose synthesis. However, whether TPS and TPP enzymes form complexes in other organisms is unknown (Bell et al., 1998). Recent structural studies of fungal TPS and TPP proteins provide insights into their active sites but do not reveal their interactions (Miao et al., 2017; Washington et al., 2024; Yang et al., 2019). In rice (*Oryza sativa*), Class II TPS proteins interact with each other and with the Class I protein OsTPS1 (Zang et al. 2011); however, the biological or enzymatic function of the rice TPS complexes remains unresolved, and whether TPSs and TPPs interact in plants has not been addressed.

Here, we show that non-catalytic maize TPS proteins form a complex with the maize TPP RA3 and with catalytic TPSs to regulate inflorescence architecture and tillering. We also show that a non-catalytic TPS enhances the enzyme activity of active TPSs and TPPs. Our studies clarify the role of non-catalytic TPSs in Tre6P metabolism and plant development.

## Results

To identify candidate *in vivo* RA3 interactors, we performed immunoprecipitation coupled with mass spectrometry (IP-MS) using extracts from ear primordia expressing a functional hemagglutinin (HA) - tagged RA3 fusion driven by its native promoter (Claeys et al., 2019). We identified ZmTPS1 as a significant RA3 interactor, with a 6-fold enrichment relative to the negative control (Fig. 1a, Table S1). Notably, RA3 itself was detected at a similar enrichment, suggesting a potential direct association between RA3 and ZmTPS1. We confirmed this interaction using bimolecular fluorescence complementation (BiFC) following transient co-expression in *Nicotiana benthamiana* leaves. We observed YFP signals in nuclei that could be resolved into speckle-like puncta by confocal microscopy, suggesting RA3 and ZmTPS1 interact in nuclei (Fig. 1b, c). A negative interaction control, using RA3 with FASCIATED EAR4 (FEA4)(Pautler et al., 2015), gave no signal (Fig. 1b).

**Figure 1.**
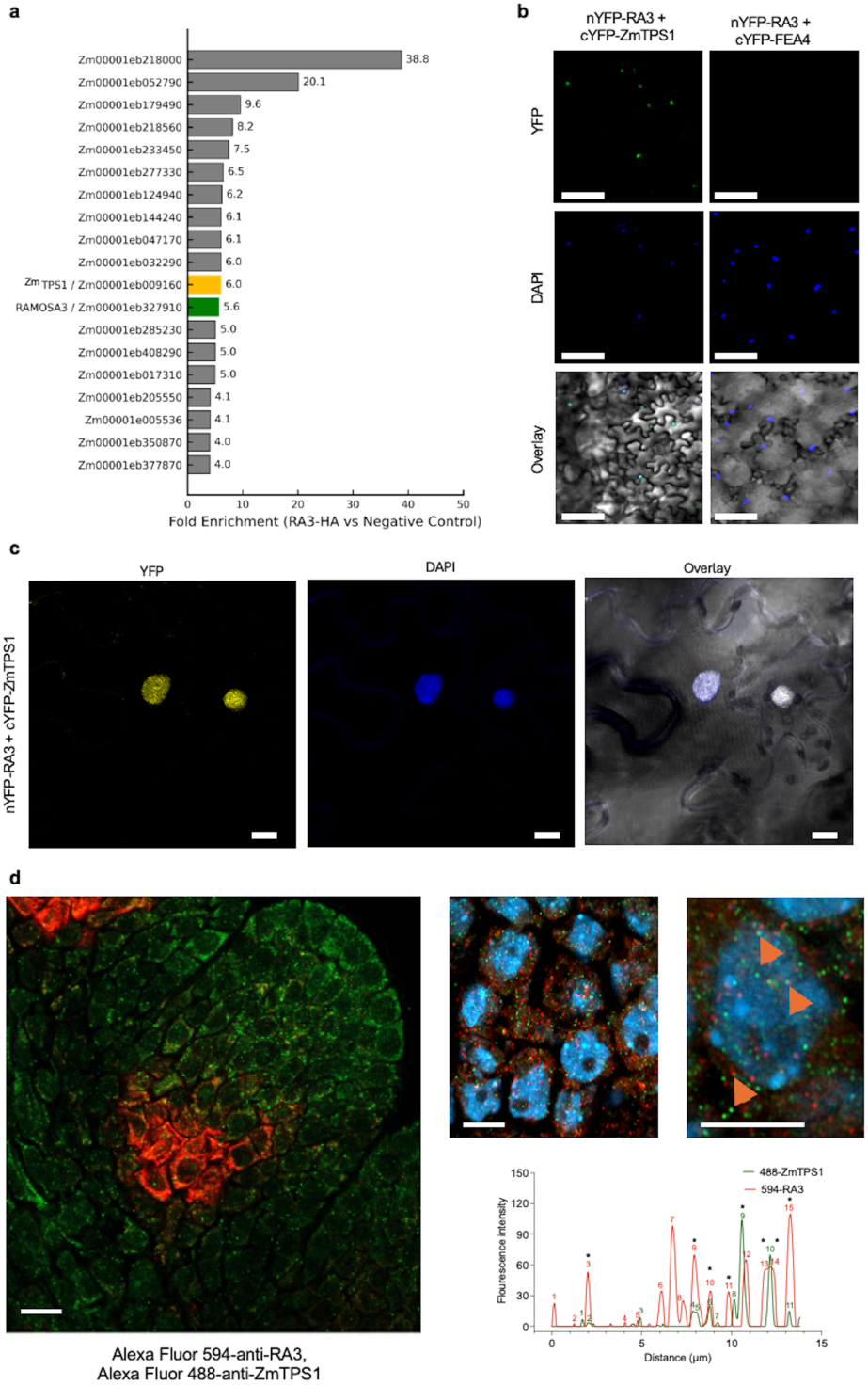
RA3 physically interacts and colocalizes with non-catalytic ZmTPS1 in maize meristems. (a) Immunoprecipitation followed by mass spectrometry (IP-MS) using HA-tagged RA3 identified ZmTPS1 (Zm00001eb099160) as a putative interactor, with ∼6-fold enrichment compared to the negative control. Other enriched proteins are shown by gene ID and fold enrichment. (b) Bimolecular fluorescence complementation (BiFC) assay in *Nicotiana benthamiana* leaves confirms interaction between RA3 and ZmTPS1. Co-expression of nYFP-RA3 and cYFP-TPS1 reconstituted YFP fluorescence, indicating a physical interaction. Nuclei were counterstained with DAPI (blue). No signal was observed with the negative control (nYFP-RA3 + cYFP-FEA4). Scale bars = 100 µm. (c) Confocal microscopy of BiFC assay shows strong YFP signals in nuclear speckles, consistent with RA3-ZmTPS1 interaction in nuclei. Scale bars = 5 µm. (d) Immunolocalization of RA3 (Alexa Fluor 594, red) and ZmTPS1 (Alexa Fluor 488, green) in developing spikelet meristems reveals colocalization in both the cytoplasm and nuclei, as indicated by merged signals and arrowheads. In enlarged panels, nuclei were counterstained with DAPI (blue). Scale bars = 10 µm. The right panels show magnified views and fluorescence intensity plots. The bottom graph quantifies the overlap of RA3 and ZmTPS1 signals (asterisks), indicating ∼24% colocalization.

We next asked if RA3 and ZmTPS1 colocalized *in vivo* in developing maize ears using immunolocalization with ZmTPS1 and RA3-specific antisera(Claeys et al., 2019). To generate ZmTPS1 specific antisera, we aligned all maize TPSs and identified a unique 143 amino acid sequence at its N-terminus (Fig. S1a). This region was tagged with a His-tag and expressed in *E. coli* then purified. Western blot analysis validated the specificity of the ZmTPS1 antisera, which recognized the His-ZmTPS1 peptide but did not cross-react with a His-ZmTPS14 peptide (Fig. S1b). ZmTPS1 signals were also absent in *zmtps1* mutant ear primordia, indicating that the ZmTPS1 antisera were specific (Fig. S1c). Next, we performed dual-label immunolocalization with ZmTPS1 and RA3-specific antisera. RA3 protein localized in cup-shaped domains at the base of axillary meristems in immature ear primordia, matching its mRNA localization (Claeys et al., 2019), while ZmTPS1 protein was more broadly localized throughout the axillary meristem. However, both proteins localized to discrete puncta, and importantly, the RA3 and ZmTPS1 puncta colocalized in the nuclei and cytoplasm of cells where their expression overlapped. We estimated the overlap to be approximately 24% (Fig. 1d).

To ask whether ZmTPS1 is a non-catalytic or catalytic TPS, we next performed a phylogenetic analysis, which grouped maize TPSs into two clades: Class I, predicted to be catalytically active, included ZmTPS11 and ZmTPS14, whereas Class II, aligning with the non-catalytic TPS-like subfamily, included ZmTPS1 and its close paralog ZmTPS12, and all other maize TPS proteins (Fig. 2a). To further evaluate the enzymatic potential of these TPSs, we aligned the amino acid residues critical for substrate binding and catalysis (Vandesteene et al. 2012; Miao et al. 2017; 2016; Washington et al. 2024) using functional TPS enzymes such as *Cryptococcus neoformans* TPS1 (CnTPS1), *E. coli* OtsA (EcOtsA), *Saccharomyces cerevisiae* TPS1 (ScTPS1), and *Arabidopsis thaliana* AtTPS1 (Fig. S2a). We focused on the conserved catalytic residues (G86, R415, K420, R453, D514, E522 in CnTPS1) and substrate stabilization and orientation residues (S84, V492, L518). These catalytic and substrate-interacting residues were conserved in maize TPS11 and TPS14, suggesting they have enzymatic activity. In contrast, ZmTPS1 contained several substitutions at these critical sites, most notably a conserved glycine residue in CnTPS1 (G86) was replaced by serine (S87) in ZmTPS1, and a conserved arginine residue in CnTPS1 (R416) was replaced by aspartate (D342) in ZmTPS1. These substitutions are predicted to disrupt catalytic activity (Fig. 2c, d), and reinforce our phylogenetic classification to support the hypothesis that ZmTPS1 and ZmTPS12 are catalytically inactive. To further confirm catalytic activities, we asked whether the different proteins could complement a *S. cerevisiae tps1* mutant (Flores et al., 2011). As expected, arabidopsis AtTPS1 and maize ZmTPS11 and ZmTPS14 could complement the yeast *tps1* mutant, promoting growth in selective media, but maize ZmTPS1 or ZmTPS12 could not (Fig.S2b).

**Figure 2.**
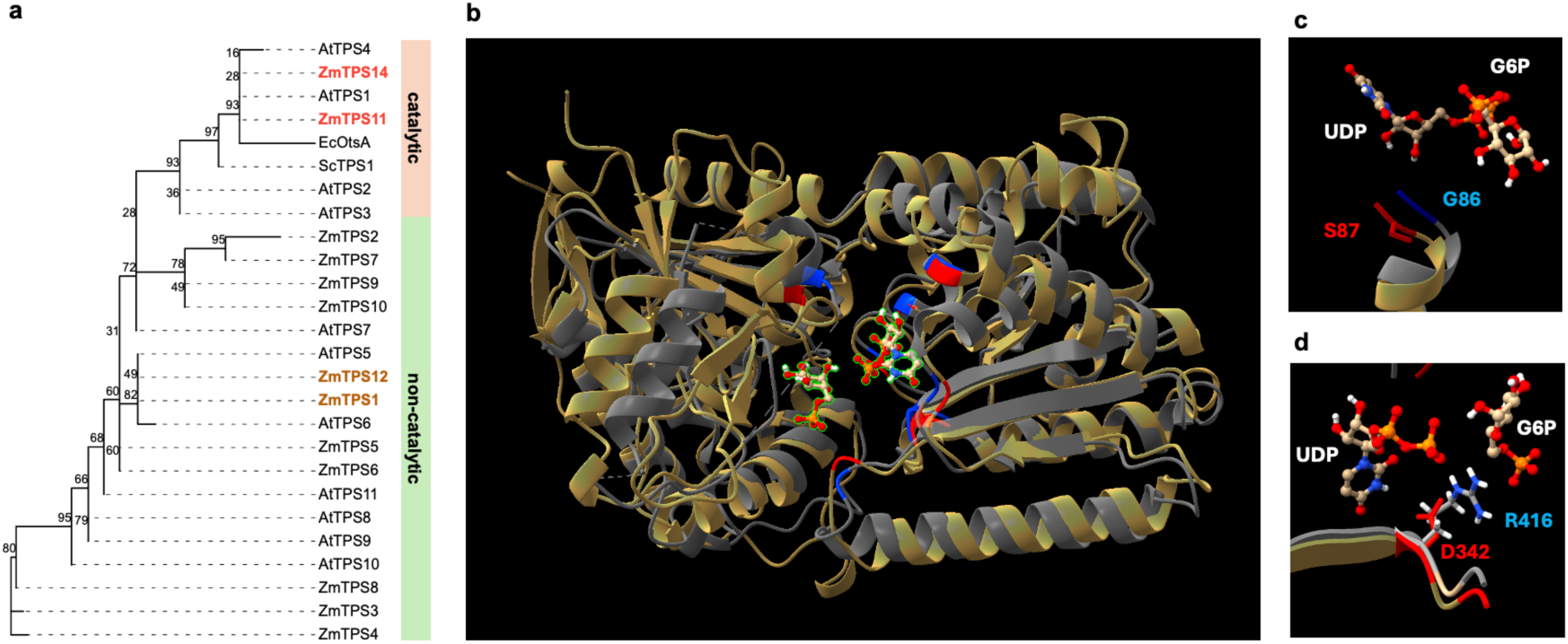
ZmTPS1 is predicted to be non-catalytic. **(a)** Phylogenetic analysis shows that ZmTPS1 and ZmTPS12 cluster with non-catalytic class II TPSs, while ZmTPS11 and ZmTPS14 group within the catalytic class I TPS clade. Bootstrap values are indicated at each node. **(b)** Structural alignment of the AlphaFold 3.0-predicted TPS domain of ZmTPS1 (gold) with the previously solved TPS domain of CnTPS1 (gray) demonstrates strong structural similarity. Residues critical for substrate binding and catalytic function in CnTPS1 are highlighted in blue, with corresponding residues in ZmTPS1 highlighted in red. Trehalose-6-phosphate and UDP are outlined in green. **(c)** Close-up view of the alignment reveals that Glycine 86 in CnTPS1 (G86, blue) is substituted by Serine 87 (S87) in ZmTPS1. This substitution at a conserved position is predicted to disrupt catalytic activity. **(d)** Close-up view of the alignment showing that Arginine in CnTPS1 (R416, blue) is substituted by aspartate 342 (D342, red) in ZmTPS1. This substitution at another conserved position is also predicted to disrupt catalytic activity.

We next asked if *ZmTPS1* mutants, as well as mutants in its close paralog *ZmTPS12*, interact genetically with *ra3*. We isolated *Mutator* transposon insertion mutants (uniform *Mu*) in coding exons of each gene, and backcrossed them twice to *ra3* (Fig. 3a). *ra3* mutants have abnormally branched ears due to indeterminate meristem growth. The single *zmtps1* or *zmtps12* mutants and the double *zmtps1;zmtps12* mutants had normal, unbranched ears. However, *zmtps1* single and *zmtps1;zmtps12* double mutants enhanced *ra3* ear branching (Fig.3b, c). Notably, the *ra3;zmtps1;zmtps2* triple mutant was strongly enhanced, with branches extending towards the middle of the ear rather than being restricted to the base, as in *ra3* single mutants (Fig. 3b). These findings indicate that ZmTPS1 and ZmTPS12 regulate ear branching together with RA3, consistent with the physical interaction we found between RA3 and ZmTPS1.

**Figure 3.**
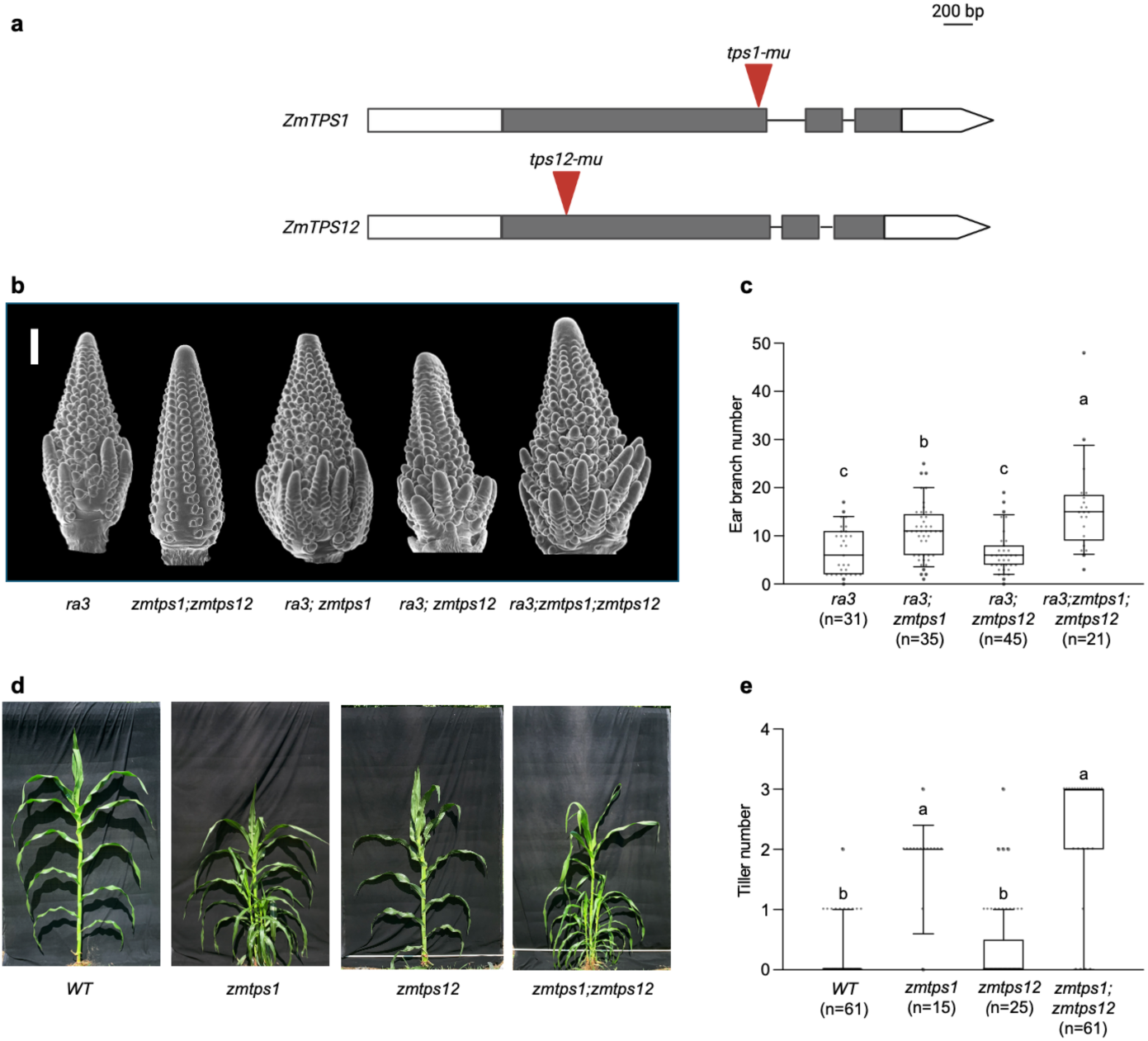
Non-catalytic ZmTPS1 and ZmTPS12 interact with RA3 to regulate inflorescence architecture and tillering in maize. **(a)** Schematic representation of ZmTPS1 and ZmTPS12 gene structures showing the positions of *Mu* transposon insertions (red arrowheads). Gray boxes indicate coding regions. **(b)** Scanning electron microscopy (SEM) images of immature ears show that zm*tps1;zmtps12* double mutants alone are phenotypically similar to wildtype ears, but when combined with *ra3*, both *ra3;zmtps1* double and *ra3;zmtps1;zmtps12* triple mutants enhanced ear branching. Scale bar = 1mm. **(c)** Quantification of ear branch number. Box plots show median, interquartile range, and outliers; letters indicate statistically significant differences (p < 0.05, ANOVA with post hoc test). **(d)** Whole-plant phenotypes of wild type WT, zm*tps1, zmtps12*, and zm*tps1;zmtps12* double mutants. Single mutant zm*tps12* appears similar to WT, while single mutant zm*tps1* and double mutants increase tiller outgrowth. **(e)** Tiller number per plant for the genotypes shown in (d). zm*tps1* single and zm*tps1;zmtps12* double mutants have significantly more tillers than WT. Statistical significance was determined as in (c).

Tre6P signaling is also associated with tiller outgrowth in maize (Dong et al., 2019), and branching in eudicots (Fichtner et al., 2021, 2017). *ra3* mutants enhance tiller formation (Klein et al., 2022), so we asked if ZmTPS1 and ZmTPS12 also regulate tillering. We counted the number of tillers in *zmtps1* or *zmtps12* single or double mutants in both wild-type and *ra3* mutant backgrounds and found enhanced tillering in *zmtps1* single mutants and in *zmtps1; zmtps12* double mutants. The tiller number was higher in the double mutants, indicating that *ZmTPS1* and *ZmTPS12* function redundantly to suppress tiller outgrowth (Fig. 3d, e). The effect of *zmtps1* and *zmtps12* mutations on tillering was also enhanced in a *ra3* background (Fig. S3a, b)

In yeast (*S. cerevisiae*), catalytic and non-catalytic TPSs form a complex (Bell et al., 1998). To investigate potential interactions between the catalytic maize TPSs 11 and 14, and non-catalytic ZmTPS1, we next performed yeast two-hybrid (Y2H) assays. Using ZmTPS1 as bait and ZmTPS11 or ZmTPS14 as a prey, we found that yeast expressing both proteins survived on selective media, indicating that ZmTPS1 has the potential to interact physically with both ZmTPS11 and ZmTPS14 (Fig.4a). Based on this interaction, and our IP-MS and genetic data, we hypothesized that ZmTPS1 might form a three-way complex with RA3 and a catalytically active TPS. We chose ZmTPS14 as a representative active TPS because its expression in ear primordia is higher than ZmTPS11 (Eveland et al., 2014). We co-expressed full-length RA3, ZmTPS1 and ZmTPS14 proteins labeled with Maltose-Binding Protein (MBP), Streptavidin (Strep), or Octahistidine (His) tags, respectively, in *Spodoptera frugiperda* (Sf9) insect cells, and performed sequential rounds of purification with an MBP column followed by a Strep column (Fig.4b). We confirmed the presence of all three proteins in the Strep column eluate by Western blotting, suggesting they form a three-component complex (Fig. 4c). However, extensive attempts using cryo-EM failed to solve the structure of the full-length protein complex, suggesting the complex was heterogeneous.

**Figure 4.**
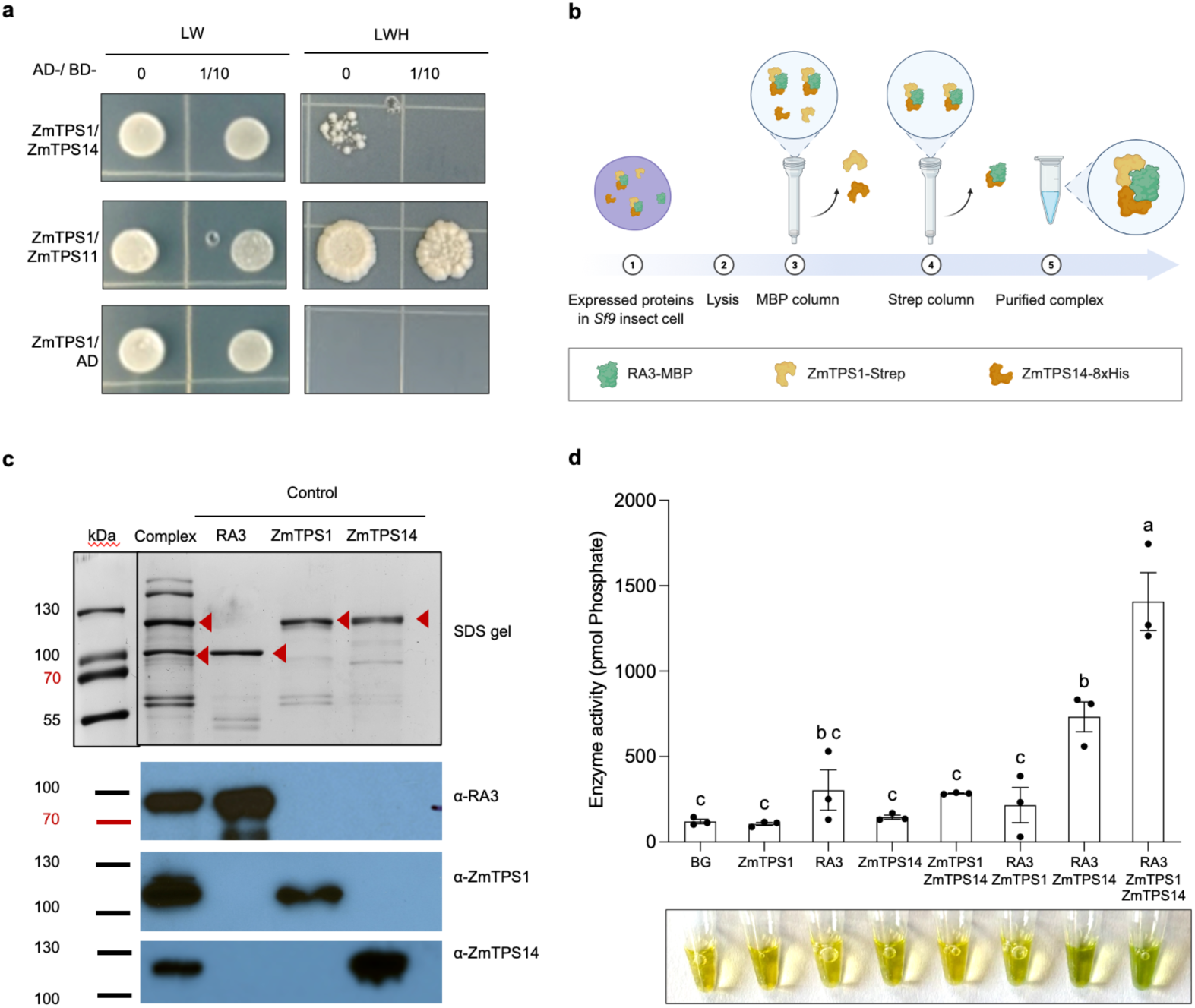
RA3, ZmTPS1, and ZmTPS14 form a functional protein complex. **(a)** Yeast-two-hybrid assays show that non-catalytic ZmTPS1 interacts with catalytic ZmTPS14 and ZmTPS11. Yeast growth on selective (-) LWH medium confirms physical interactions between protein pairs. **(b)** Schematic diagram of the two-step purification strategy for isolating the RA3-ZmTPS1-ZmTPS14 complex. Full-length RA3, ZmTPS1, and ZmTPS14 were co-expressed in insect cells, each with a unique affinity tag (MBP, Strep, and His, respectively), and purified sequentially via MBP and Strep affinity columns. **(c)** SDS-PAGE and Western blot analysis of the purified protein complex. Each protein was detected using specific antisera, red arrows mark bands corresponding to RA3, ZmTPS1, and ZmTPS14 within the complex. **(d)** Coupled enzymatic activity assays show that ZmTPS1 enhances the catalytic activity of RA3-ZmTPS14, while ZmTPS1 alone is non-catalytic. Bars represent mean ± SD; different letters denote statistically significant differences (one-way ANOVA with Tukey’s test, p < 0.05). Lower panel: reaction tubes from the enzyme assay corresponding to each test condition.

We next asked whether the non-catalytic ZmTPS1 protein might affect the catalytic activity of ZmTPS14 and RA3. We expressed and purified Maltose binding protein (MBP) tagged RA3 from *E. coli*, and Strep-tagged ZmTPS1 and ZmTPS14 from Sf9 insect cells. After affinity purification of each protein, we used *in vitro* assays to measure the combined activity of ZmTPS1, ZmTPS14 and RA3. We developed a coupled TPS-TPP assay, using the TPS substrates UDP-glucose and glucose-6-phosphate, and measuring phosphate released by TPP activity. As expected, neither of the three proteins alone exhibited significant activity above background in this coupled assay (Fig. 4d). As expected, the active TPS, ZmTPS14, but not ZmTPS1, showed significant activity when combined with RA3. However, when we combined ZmTPS1, RA3 and ZmTPS14, we observed a significant increase in enzymatic activity, despite ZmTPS1 having no activity alone or in combination with RA3 (Fig. 4d). These results indicate that ZmTPS1, despite having no detectable enzymatic activity itself, enhances the coupled enzymatic activities of ZmTPS14 and RA3. We next used AlphaFold 3 (Abramson et al., 2024) to model the potential interactions of ZmTPS1, ZmTPS14, and RA3, and the three proteins were predicted to form a heterotrimer. However, the interface predicted template modeling (ipTM) score was moderate. We also asked whether the TPS domains in ZmTPS1 and 14, might promote complex formation. The two TPS domains were predicted to interact strongly, with an ipTM score of 0.89, suggesting that the two TPS domains of ZmTPS1 and ZmTPS14 stimulate the formation of a complex (Fig. 5).

**Figure 5.**
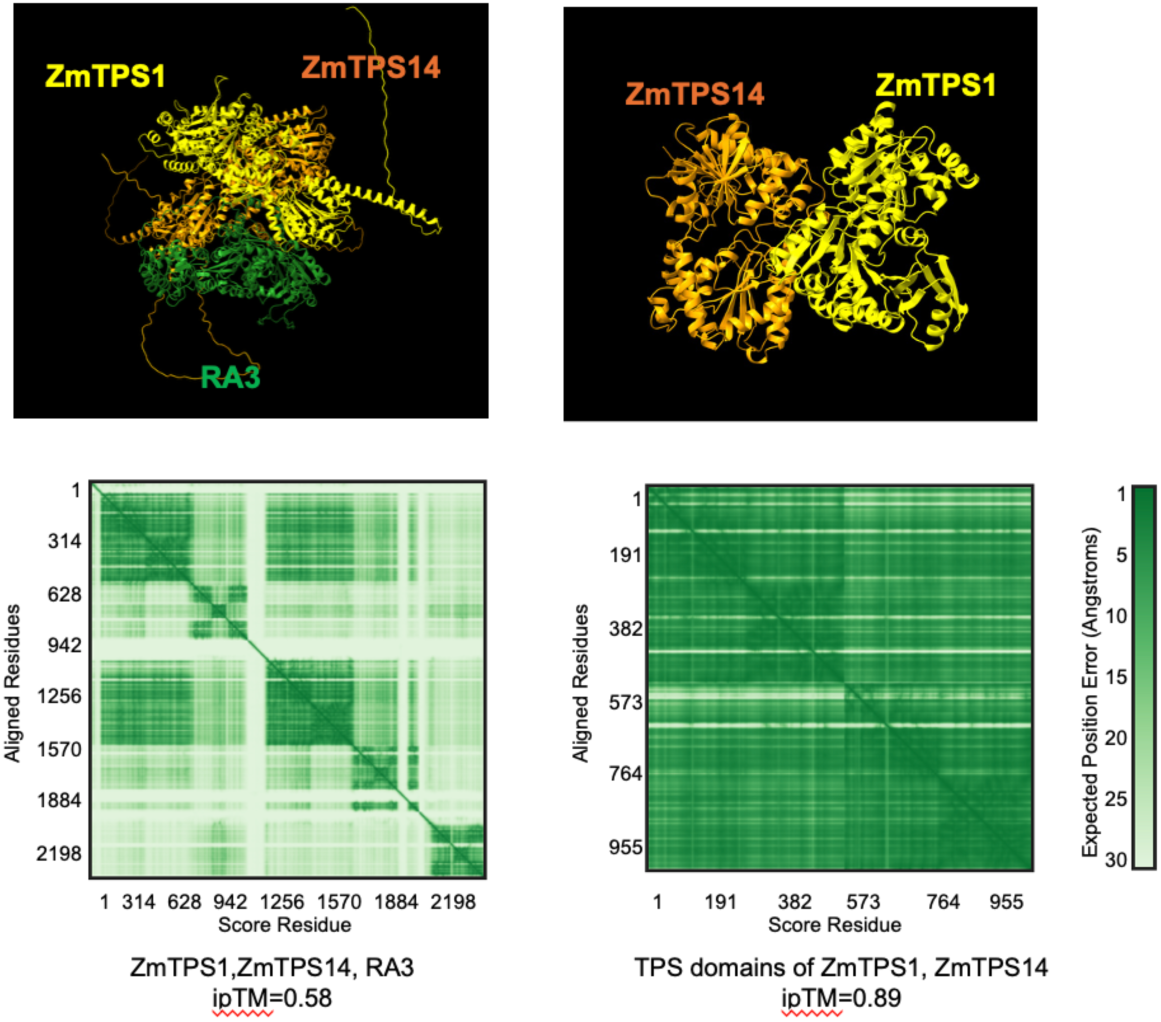
The TPS domains of ZmTPS1 and ZmTPS14 are predicted to form a complex. AlphaFold 3.0 structural predictions **Upper panels:** Predicted 3D protein structure of the RA3-ZmTPS1-ZmTPS14 complex (left) and TPS domains only (right). **Lower panels.** Predicted aligned error (PAE) matrices indicating predicted interfaces between proteins. The score was high for the ZmTPS1-ZmTPS14 TPS domain interaction (0.89) and moderate for the full-length complex (0.58), supporting a model where TPS domain-mediated interactions stabilize the complex.

To understand the functions of catalytically active TPSs in maize growth and development, we isolated a *zmtps11* Mu-insertion line (Fig. 6a) and generated a *zmtps14* CRISPR-Cas9 knockout line (Fig. 6a). We generated two independent CRISPR-Cas9-induced alleles of the *ZmTPS14* gene. The first allele contained a G deletion in exon 1, and the second carried a T insertion in exon 2. Both mutations resulted in a frameshift and a premature stop codon, predicted to disrupt ZmTPS14 function. We selected the T insertion allele for further phenotypic analysis. The single *zmtps11* or *zmtps14* mutant embryos developed normally, with clearly visible shoot apical meristem and leaf primordia after clearing (Fig. 6c). In contrast, *zmtps11; zmtps14* double mutant embryos arrested early, with abnormal shoot meristems, and did not develop further. This arrest aligns with the shift from pattern formation to organ expansion (Sheridan and Clark, 1993) (Fig. 6 b, c). The *zmtps11; zmtps14* double mutant kernels were also smaller and failed to germinate (Fig 6b). To ask if the developmental defects in the double mutants were caused by a lack of Tre6P, we performed embryo rescue experiments using media supplemented with 2mM Tre6P, 100 mM Trehalose, or 100 mM sucrose. (Fig. 6d). The double mutant embryos were partially rescued on media containing a membrane-permeable Tre6P analog (Griffiths et al., 2016), but not by trehalose or sucrose. These observations support the notion that ZmTPS11 and ZmTPS14 are catalytically active TPSs, acting redundantly to catalyze Tre6P and promote maize embryo and endosperm development.

**Figure 6.**
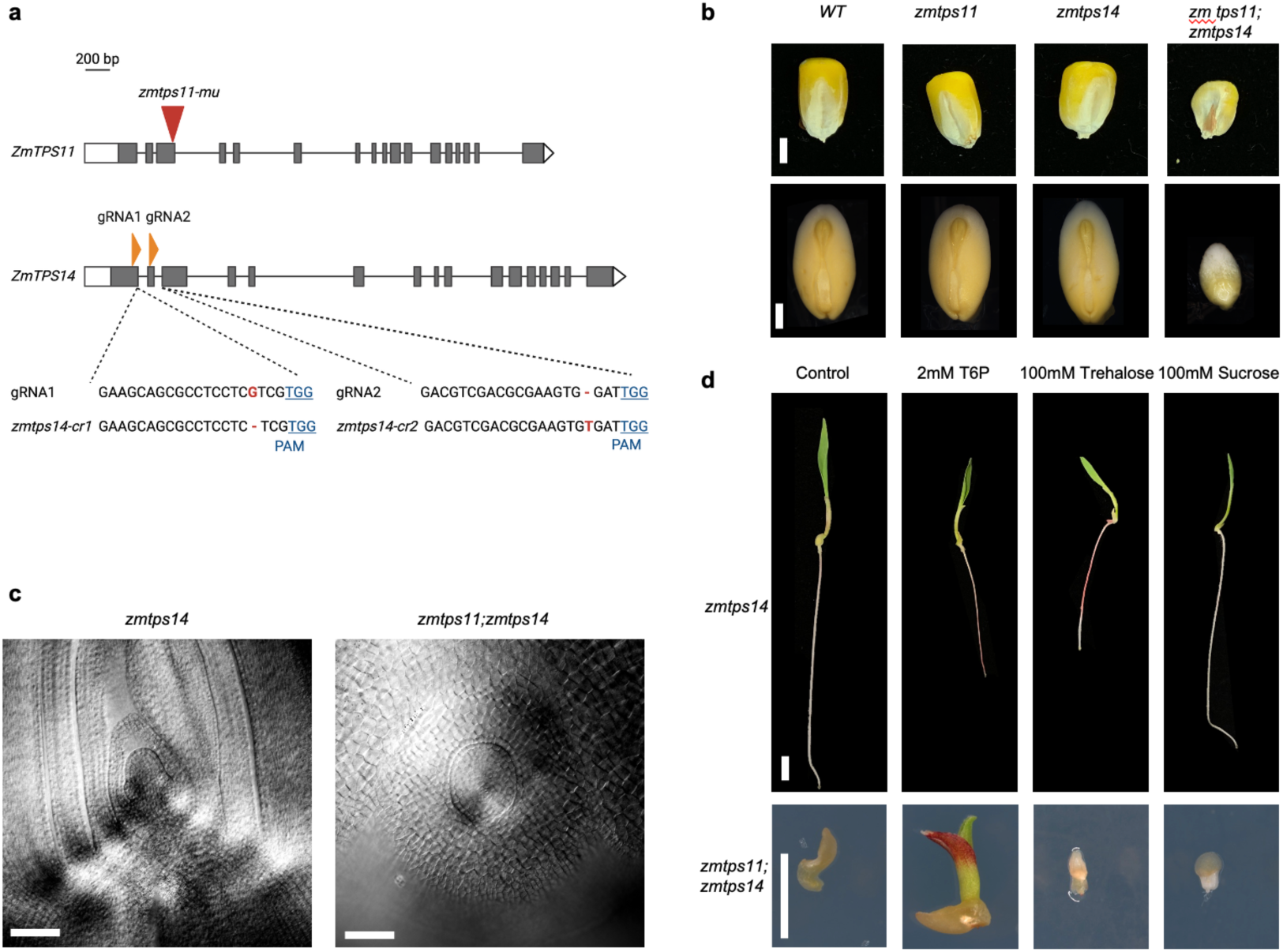
ZmTPS11 and ZmTPS14 are required for maize embryo and kernel development. (a) Gene models of *ZmTPS11* and *ZmTPS14,* showing the zm*tps11-Mu* insertion site and CRISPR guide RNA target sites (gRNA1 and gRNA2) for *ZmTPS14*. Target sequences and corresponding PAM sites of ZmTPS14 are indicated. (b) Mature kernels (top) and dissected embryos (bottom) of wild type, *zmtps11*, *zmtps14*, and zm*tps11; zmtps14*. Single mutants develop normally, while double mutants have severe reductions in kernel and embryo size. (c) Microscopic images of zm*tps14* and zm*tps11; zmtps14* embryos at 21 DAP. *zmtps14* embryos develop normally. In contrast, zm*tps11; zmtps14* double mutant embryos arrest with a coleoptile-enclosed meristem and no further organ development. (d) Germination rescue assay of *zmtps14* and *zmtps11;zmtps14* embryos cultured on media containing 2 mM Tre6P, 100 mM trehalose, or 100 mM sucrose. Only Tre6P partially rescues the growth of *zmtps11;zmtps14* embryos, supporting a specific requirement for Tre6P in embryo development. Scale bars: 2 mm (b), 10 µm (c), 1cm (d, top), 5 mm (d, bottom).

## Discussion

The branching architecture of maize ears is tightly regulated by complex genetic and metabolic networks that coordinate meristem fate, energy signaling, and developmental transitions (Lindsay et al., 2024). Tre6P is an important developmental and physiological signal in plants, and although plant genomes encode a small family of TPSs, many of them have no enzymatic activity, raising questions about their biological roles. Our study defines distinct yet cooperative roles for both catalytic and non-catalytic maize TPS proteins. Notably, we found that non-catalytic maize TPS1 and TPS12 regulate inflorescence determinacy and axillary bud dormancy in coordination with RA3. We also found that non-catalytic TPSs interact physically with both RA3 and catalytic TPSs to control Tre6P and trehalose biosynthesis.

We identified ZmTPS1 as an RA3-associated protein through IP-MS analysis, suggesting a physical and potentially functional interaction between these two proteins. Further studies revealed that ZmTPS1 and RA3 colocalize in cytoplasmic and nuclear puncta at the base of developing spikelet meristems, consistent with a hypothetical moonlighting role of RA3 in transcriptional regulation (Claeys et al., 2019; Demesa-Arevalo et al., 2021). Single and double mutants of *zmtps1* and *zmtps12* lacked any obvious phenotypes. However, in a *ra3* mutant background, the loss of *zmtps1* and *zmtps12* dramatically enhanced ear branching, with triple mutants having more branches than double or single mutants. Given that ZmTPS1 and ZmTPS12 are non-catalytic, their genetic interaction with RA3 suggests that they function as regulatory factors rather than as enzymes.

Although multimeric TPS-TPP complexes have long been proposed (Bell et al., 1998; Chen et al., 2021; Zang et al., 2011), structural studies have focused mainly on single subunits. To explore the intersection of regulatory and enzymatic functions, we reconstituted a RA3-ZmTPS1-ZmTPS14 complex in insect cells, mimicking the yeast trehalose synthase complex model outlined by Bell et al. (1998). Although ZmTPS1 is catalytically inactive, it retains key substrate-binding residues and differs from the active ZmTPS14 by only a conservative V492L substitution in the uracil-binding site (Miao et al. 2017; Washington et al. 2024), a change unlikely to affect substrate recognition. In a coupled enzyme assay, ZmTPS1 enhanced the enzymatic activity of RA3 and ZmTPS14, suggesting that it facilitates substrate positioning or stabilizes the active complex, despite lacking catalytic function. In addition, AlphaFold modeling of RA3-ZmTPS1-ZmTPS14 and the identification of conserved amino residues important for complex formation in ZmTPS1 and ZmTPS14 support the formation of higher-order assemblies.

We also uncovered roles of non-catalytic ZmTPS1 and ZmTPS12 in repressing tiller bud outgrowth. The enhanced tillering observed in both *zmtps1 zmtps12* and *ra3* mutant backgrounds suggests that RA3 and class II TPS proteins act in the same genetic pathway. The transcription factors GRASSY TILLERS1 (GT1) and TEOSINTE BRANCHED1 (TB1), known repressors of bud outgrowth, positively regulate the expression of class I and II TPS genes, as shown by their downregulation in *gt1* and *tb1* mutants (Dong et al., 2019). Intriguingly, RA3 expression is elevated in *gt1*, and TB1 directly binds to the RA3 promoter (Dong et al., 2019), indicating that RA3 is negatively regulated by GT1 and may be transcriptionally modulated by TB1. Together, our findings support a model in which GT1 and TB1 control bud dormancy and tillering by regulating the expression of RA3 and catalytic and non-catalytic TPS proteins. Through this regulation, Tre6P signaling is positioned as an integrator of developmental and metabolic signals controlling bud outgrowth.

We also identified essential roles for catalytic ZmTPS11 and ZmTPS14 in maize embryo development. *zmtps11;zmtps14* double mutants arrested early in development, closely resembling the stage-specific embryo block observed in arabidopsis *tps1* mutants (Eastmond et al., 2002a). Notably, this defect could be partially rescued by low concentrations of exogenous Tre6P, but not by trehalose or sucrose, suggesting that Tre6P functions primarily as a signaling molecule in this context, rather than a metabolite. These findings demonstrate a conserved role for catalytic TPS enzymes in embryo and seed viability and underscore the broader importance of Tre6P as a metabolic signal that integrates sugar availability with developmental progressions. Collectively, our findings support a model in which both catalytic and non-catalytic TPS proteins, along with a TPP protein, such as RA3, assemble into regulatory complexes that integrate sugar signaling with developmental control. Our findings expand the classical view of TPS proteins, demonstrating that catalytically inactive family members function as scaffolds and signaling mediators, revealing a previously overlooked aspect of metabolic regulation in plant architecture.

## Author contributions

Thu M. Tran: Conceptualization; performed research and co-wrote the manuscript. Hannes Claeys: generated *zmtps14-crispr* lines, performed yeast complementation and yeast-two-hybrid experiments, expressed RA3 protein, and isolated *zmtps11-mu*. María Jazmín Abraham-Juárez: developed ZmTPS1 and ZmTPS14 antibodies. Xiaosa Xu: performed BIFC assays. Kevin Michalski: assisted and advised on the expression of ZmTPS1, ZmTPS14, and RA3 in insect cells. Tsung Han Chou: assisted and advised with the expression of TPS domains of ZmTPS1 and ZmTPS14 in insect cells. Sessen D. Iohannes: processed immunolocalization data and assisted with phenotyping mutants. Panagiotis Boumpas: performed immunolocalizations and embryo rescue. Z’Dhanne Williams: genotyped mutants. Samatha Sheppard: genotyped mutants and performed enzyme assays. Son L. Vi: performed RA3-HA IP-MS and isolated *zmtps1-mu* and *zmtps12-mu*. Matthew J. Paul and Cara Griffiths: provided DMNB-Tre6P. Hiro Furukawa: advised on protein expression and analysis; David Jackson: conceptualization and co-wrote the manuscript.

## Acknowledgments

The study was funded by the NSF award IOS-2131631. The design, synthesis and experimental testing of DMNB-T6P was funded by UKRI-BBSRC Selective Chemical Intervention in Biological Systems (BB/D006112/1) to Matthew Paul and Ben Davis, and Matthew Paul and Cara Griffiths acknowledge the Delivering Sustainable Wheat Institute Strategic Program (BB/X011003/1). We thank the Maize Genetics Cooperation Stock Center for providing Uniform Mu mutant lines. We the Pedmale lab for the use of the confocal microscope and Joshua-Tor laboratory for the use of mass photometry equipment. We thank the Martienssen lab and Joseph Simorowski for the use of the ChemiDoc Touch Imaging System (Bio-Rad) and fluorescence microscope. We thank the Mass Spectrometry Facility at Cold Spring Harbor Laboratory for sample processing. We thank the Uplands farm team, including Tim Mulligan, Kyle Schlecht, and Autumn Harrison, for plant care. We thank Patrick Van Dijck for sharing the YSH567 yeast strain. and our dedicated summer students who assisted with field and lab work: Emma Morley, Spencer Elliott, Jessica Finger, Taylor Petrizzo, Matt Venezia, and Kylie Chen.

## Methods

### Plant Materials and Growth Conditions

The *ra3-fea1* (Satoh-Nagasawa et al., 2006), *zmtps1-mu* (Mu1055231)*, zmtps11-mu* (Mu1030345) (McCarty et al., 2013), and *zmtps12-mu* (IlluminaMu 393550) (Williams-Carrier et al., 2010) alleles were isolated from Mutator transposon lines and backcrossed to B73 or *ra*3 in the B73 background.

To generate CRISPR-Cas9-knockout mutants, guide RNAs (gRNAs) targeting ZmTPS11 and ZmTPS14 were designed using CRISPR-P (http://cbi.hzau.edu.cn/crispr/), and an array containing two gRNAs expressed from independent maize U6-derived promoters was synthesized, with gRNAs GAAGCAGCGCCTCCTCGTCG and GACGTCGACGCGAAGTGGAT. This array was cloned into the pMCG1005-Cas9 binary vector and transformed into the Hi-II background. Mutations were identified by Sanger sequencing and backcrossed at least three times to *zmtps1, zmtps12, ra3,* or *zmtps11* (B73) before phenotyping. The inserted T-DNA containing Cas9 and the gRNA array was eliminated after the first backcross to ensure the stability of mutations. The primers for cloning are listed in Table S2.

Plants for crossing and phenotyping were genotyped (primers are listed in Table S2).

### Bimolecular Fluorescence Complementation (BiFC) Assay

A BIFC assay was used to detect interaction between RA3 and ZmTPS1 proteins in *Nicotiana benthamiana* leaves. Coding sequences of RA3 and ZmTPS1 were cloned to generate fusions with the N-terminal (nYFP) and C-terminal (cYFP) fragments of YFP, respectively, as previously described (Li et al., 2009). *Agrobacterium tumefaciens* strain GV3101 carrying the constructs was infiltrated into 4-week-old *N. benthamiana* leaves at OD= 0.01 using a syringe. Plants were maintained under standard growth conditions for 4 days post-infiltration. YFP fluorescence, indicating protein–protein interaction, was detected using confocal laser scanning microscopy (excitation 514 nm, emission 525–550 nm). Leaves co-infiltrated with RA3 and FEA4 fusions with the N-terminal (nYFP) and C-terminal (cYFP) served as negative controls. At least three biological replicates were imaged per condition. Fluorescence intensity and localization were quantified using ImageJ. The primers for cloning are listed in Table S2.

### Phenotyping

For ear and tiller phenotyping, plants were grown in field locations in Cold Spring Harbor, NY, USA, or Lloyd Harbor, NY, USA (June–September), or Hawaii (November–February) between 2020-2025. Ears were collected after anthesis to count the branches. Developing ears were dissected around 6 weeks after planting for scanning electron microscopy and mounted on stubs to image on a JEOL Benchtop Scanning Electron Microscope. All phenotyping experiments were performed on segregating populations, and each plant was individually genotyped.

### Phylogenetic analysis

The TPS protein sequences from *Cryptococcus neoformans*, *Escherichia coli*, *Saccharomyces cerevisiae*, *Arabidopsis thaliana*, and *Zea mays* were retrieved from public databases. The published amino acid residues critical for substrate binding and catalysis were aligned using MUSCLE implemented in MEGA11 (Tamura et al., 2021). A maximum likelihood phylogenetic tree was constructed in MEGA11 with 1,000 bootstrap replicates to assess node support. The final tree was exported and visualized using the Interactive Tree of Life (iTOL) web tool (Letunic and Bork, 2021).

### Embryo phenotypes

Maize embryos segregating *zmtps14* and *zmtps14;zmtps11* on the same ear were dissected at 21 days after pollination. Embryos were fixed in FAA (10% formalin, 5% acetic acid, and 45% ethanol) and cleared in methyl salicylate. Embryos were observed by light microscopy with at least ten embryos per genotype. For embryo rescue, immature embryos (14 days after pollination) from bleach-sterilized ears segregating zm*tps11*, *zmtps14,* and *zmtps11; zmtps14* were excised under sterile conditions using fine forceps and a scalpel. The endosperm was kept for DNA extraction and genotyping. Embryos were placed embryo-side up on solid Murashige and Skoog (MS) medium, with the addition of 2mM of membrane-permeable Tre6P precursor, DMNB-Tre6P dissolved in DMSO (Griffiths et al., 2016), or 100 mM Trehalose, or 100 mM Sucrose. Plates were sealed with micropore tape and incubated in a growth chamber at 25 °C under a 16 h light/8 h dark cycle. Germination was observed after 7 days, and embryos were then used to extract DNA and perform PCR to confirm genotypes.

### Immunoprecipitation coupled with mass spectrometry (IP-MS)

The RA3-HA transgene was constructed by amplifying genomic fragments and fusing the HA sequence in frame at the C terminus. Confirmed clones were transferred to *Agrobacterium* and transformed into maize, as described (Krishnakumar et al., 2015). The positive plants were then backcrossed to B73 and to the *branched silkless1;tunicate1* double mutant background (Chuck et al., 2002; Han et al., 2012) to increase the amount of meristem tissue.

Frozen ear primordium tissue was ground in liquid nitrogen and extracted in IP buffer (50 mM Tris-HCl (pH 7.5), 150 mM NaCl, 0.5% Triton X-100, 1 mM EDTA) with protease inhibitors. Lysates were rotated for 5 min, filtered, sonicated, and clarified by centrifugation (2 × 10 min, ≥16,000 ×g). Anti-HA antibody (Abcam ab9110; 1 µg per 10 µL Dynabeads Protein A) was bound to the beads and incubated with lysate for 30–60 min at 4 °C. The beads were then washed 4 times in IP buffer (the first two washes with half-strength inhibitors, the final two in fresh tubes without inhibitors). The beads were snap-frozen in liquid nitrogen and stored at –80 °C before MS analysis. The samples were then analyzed by the Mass Spectrometry Facility at Cold Spring Harbor Laboratory, following standard protocols.

### Generation and Validation of TPS and RA3 Antibodies

Antibodies against RA3 were generated in rabbits using a peptide produced in *E. coli* consisting of the first 80 residues of RA3 fused to a His-tag as the antigen and purified with the same peptide as described (Claeys et al., 2019). Antibodies against ZmTPS1 or ZmTPS14 were generated in guinea pigs using specific regions of ZmTPS14 or ZmTPS1 fused to a His-tag as the antigen and purified with the same peptides (Fig.S1b). Antibody specificity was validated against 1µg each of purified recombinant proteins: His-ZmTPS1; His-ZmTPS14. After SDS-PAGE and staining, proteins were transferred to a PVDF membrane by semi-dry transfer, followed by blocking with 5% non-fat milk. Blots were incubated with primary antibody (1:3000 dilution) and horseradish peroxidase (HRP)-conjugated anti-guinea pig secondary antibody (1:6000 dilution), and visualized using Clarity Western ECL Substrate (Bio-Rad) with a 5-second exposure (Fig.S1b).

### Immunolocalizations

Immunolocalization was performed on vibratome sections of 2-3 mm ears harvested from 6-week-old B73 or zm*tps1* plants as described (Tran et al., 2021). RA3 was detected using the purified primary antibody and Alexa Fluor Plus 594 coupled Goat anti-Rabbit IgG (H+L) (Invitrogen, cat. no. A32740) secondary antibody. ZmTPS1 and ZmTPS14 were detected using the purified primary antibody and Alexa Fluor 488-Goat anti-Guinea Pig IgG (H+L) secondary antibody (Invitrogen, cat. no. A-11073). Nuclei were counterstained using DAPI (Invitrogen). Imaging was performed on a Zeiss LSM 900 confocal microscope.

### Image analysis

For analyzing the co-localization of ZmTPS1 and RA3 signals, the fluorescence intensity of Alexa Fluor 488 and Alexa Fluor Plus 594 images were adjusted so that the fluorescence intensity of the empty regions was close to zero. The images were then denoised using the DEspeckle tool (Schneider et al., 2012). The Plot Profile tool (Schneider et al., 2012) quantified the fluorescence intensities of Alexa Fluor 488 and Alexa Fluor Plus 594 within the same cell. The total number of peaks in the Alexa Fluor 488 signal was labeled as the number of ZmTPS1 peaks, while the total number of peaks in the Alexa Fluor Plus 594 signal was labeled as the number of RA3 peaks, with overlapping peaks identified as co-localization of the two proteins. Ten cells were analyzed.

### Yeast complementation and Yeast-2-Hybrid

For yeast complementation, the coding sequences of AtTPS1, ZmTPS1, ZmTPS11, ZmTPS12, or ZmTPS14 were cloned into the pYX212 plasmid for constitutive expression under the glyceraldehyde 3-phosphate dehydrogenase (GPD) promoter (Miller, 1998). The primers for cloning are listed in Table S2. All sequences were verified by Sanger sequencing before transformation into the *S. cerevisiae* YSH567 *tps1; tps2* strain (W303-1A, *tps1 ::TRP1, tps2 ::LEU2*) (Hohmann et al., 1993). Cells were grown in liquid culture in SD-Ura medium to an OD600 of 1, spotted on SD-Ura containing 2% galactose (control) or 2%glucose (Sigma-Aldrich), and grown at 30 °C for 2 days. Expression of TPS-HA fusion proteins was assessed by western blot using monoclonal anti-HA antibody from mouse (Sigma-Aldrich).

For the yeast-2-hybrid, the coding sequence of ZmTPS1 was cloned into the GAL4 DNA-binding domain vector pGBKT7, while ZmTPS11 and ZmTPS14 were cloned into the GAL4 activation domain vector pGADT7. The primers for cloning are listed in Table S2. Yeast two-hybrid interaction assays were performed using the Matchmaker Gold system (Clontech). Plasmids were co-transformed into the *Saccharomyces cerevisiae* strain AH109 following the lithium acetate method. LW (–Leu/–Trp) medium selects for yeast containing both the activation domain (AD) and binding domain (BD) plasmids. Growth on LWH (–Leu/–Trp/–His) medium indicates a positive protein-protein interaction through activation of the HIS3 reporter gene.

### Protein expression and purification

For enzyme assays, RA3 coding sequences were cloned into pMAL-c5x (New England Biolabs). The primers for cloning are listed in Table S2. All sequences were verified by Sanger sequencing before transformation into the Rosetta *E. coli* strain. Cultures were grown to an optical density of 600 nm (OD600) of 0.6 at 37 °C, then cooled to 16 °C before the addition of isopropyl-β-d-thiogalactopyranoside to a final concentration of 0.5 mM. The cultures were further grown for an additional 12–16 h at 16 °C. Culture harvesting and purification of MBP-RA3 was performed with the pMAL expression system (New England Biolabs) according to the manufacturer’s instructions.

Insect codon-optimized coding sequences of ZmTPS1 and ZmTPS14 were cloned into the pFW plasmid (Podbielski et al., 1996), and proteins were expressed in Sf9 insect cells (Thermofisher, 11496015) using the EarlyBac expression method (Furukawa et al., 2021). Sf9 cells were grown in CCM3 media (Cytiva) to a density of ∼4 × 10^6^ cells/ml, then infected with freshly collected P1 virus (30 ml virus per 1 l cells), and incubated for 48h at 27 °C with shaking. Cells were collected by centrifugation at 4000g for 10 min at 4 °C, washed once in Tris-buffered saline (TBS; 20 mM Tris, 150 mM NaCl, pH 7.5), and frozen at −80 °C as cell pellets. Cells were thawed and suspended in Hepes Buffered Saline (HBS, 20 mM HEPES pH 7.5, 150 mM NaCl) with 1 mM phenylmethylsulfonyl fluoride and lysed by two passes through an Avestin cell disruptor operating at ∼10,000 psi. The lysate was cleared by ultracentrifugation at 40,000 rpm for 30 min at 4 °C in a Ti-45 rotor (Beckman Coulter), and the supernatant was passed through a 5 ml Strep-Tactin Sepharose column (IBA Lifesciences). The column was washed with 10 volumes of HBS before elution with the same buffer supplemented with 3 mM Desthiobiotin (Sigma).

For co-expression, ZmTPS1, ZmTPS14, and RA3 sequences were cloned into the pFW expression vector, linked by a P2A self-cleaving peptide under a single promoter (Kim et al., 2011). The construct was expressed in Sf9 insect cells. Next, the protein mixture underwent sequential rounds of purification for the MBP tag and Strep tag. The presence of all three proteins was confirmed by western blot using the antibodies raised against each individual protein. Protein purity and concentration were evaluated by SDS–PAGE, Nanodrop spectrophotometry, and size-exclusion chromatography (SEC) using a Superose 6 Increase column (GE Healthcare) in an HBS buffer. The final complex was concentrated to at least 1 mg/mL using an Amicon Ultra centrifugal filter unit, 10 kDa MWCO.

### Western blotting

0.5 µg of each purified recombinant protein was run in an 11% SDS-PAGE gel and then transferred to a PVDF membrane by semi-dry transfer, followed by blocking with 5% non-fat milk. Blots were incubated with primary antibody (1:500 dilution) at 4°C overnight. The blots were then incubated with HRP-conjugated anti-guinea pig or anti-rabbit IgG (1:2000, Sigma-Aldrich) for one hour at room temperature and developed using enhanced chemiluminescence (ECL) substrate (Bio-Rad) and visualized using ChemiDoc Touch Imaging System (Bio-Rad) with a 5-10 second exposure.

### AlphaFold Structural Modeling

Protein structure and interaction modeling were performed using AlphaFold 3 (Abramson et al., 2024). Full-length sequences of ZmTPS1, ZmTPS14, and RA3 were submitted individually and in pairwise or trimeric combinations using the multimer mode. Domain-specific models were also generated by isolating the annotated TPS and TPP domains based on Pfam predictions. Interface predicted template modeling (ipTM) scores were used to evaluate the likelihood and confidence of inter-protein interactions. All predictions were run with default parameters unless otherwise specified.

### Enzyme assays

For *in vitro* assays, 40 µmol of each purified protein was incubated for 30 min at 28 °C in buffer containing 10 mM Tris-HCl, 5 mM MgCl_2_, 0.1 mg/ml BSA, and 10 mM glucose 6-phosphate, UDP-glucose (Sigma-Aldrich) in a final volume of 25µl. Phosphate release was measured using a colorimetric assay at OD600 in a Synergy H4 Hybrid Reader using the Serine/Threonine Phosphatase Assay System (Promega). The activity was expressed as the free Pi released after reactions. The final enzyme activity was calculated by subtracting the background, and all samples were run in 3 independent replicates.

### Data analysis

Statistical analyses were performed using GraphPad Prism (version 9), applying appropriate tests as indicated in the figure legends. Schematic diagrams in Figs. 2a, 3b, and 5a were created using Biorender.

---

**Figure S1.**
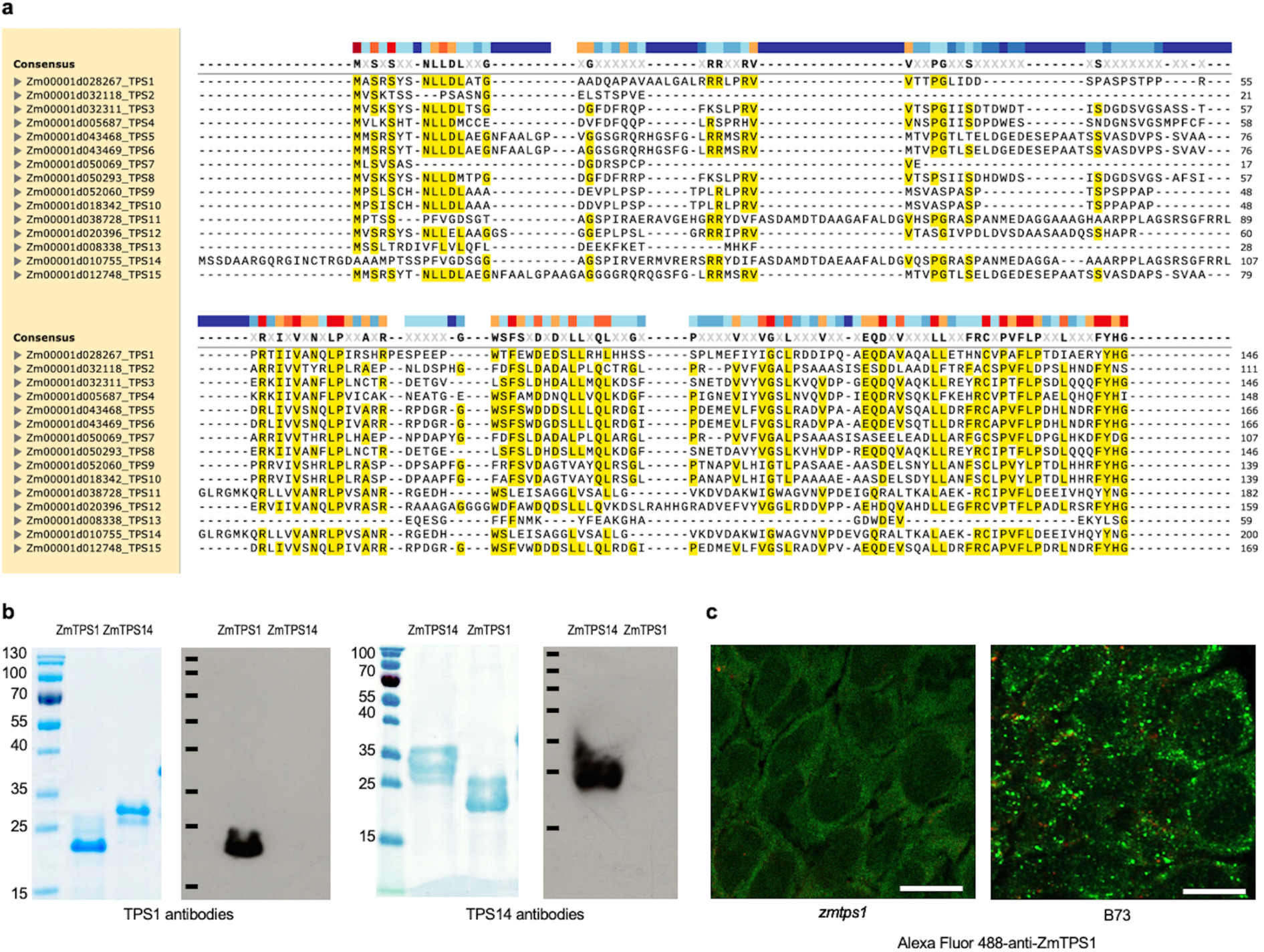
Alignment of maize TPS sequences and validation of TPS antisera specificity. (a) Amino acid alignment of the N-terminal regions of maize TPS proteins reveals a less conserved 143-residue segment in ZmTPS1 and a ∼200-residue segment in ZmTPS14. These regions were used to generate specific antisera. (b) Left: Coomassie-stained SDS-PAGE gels of purified His-tagged recombinant proteins (His-TPS1, His-TPS14. Right: western blots show that anti-ZmTPS1 and anti-ZmTPS14 antisera selectively detect their corresponding targets, with no cross-reactivity. (c) Immunolocalization negative control in a *tps1* mutant spikelet shows a lack of ZmTPS1 signal (left) and a B73 ear shows ZmTPS1 signal in green puncta (right). Samples were stained with anti-ZmTPS1 followed by a secondary antisera coupled to Alexa Fluor 488 (green). Scale bar = 10 µm.

**Figure S2.**
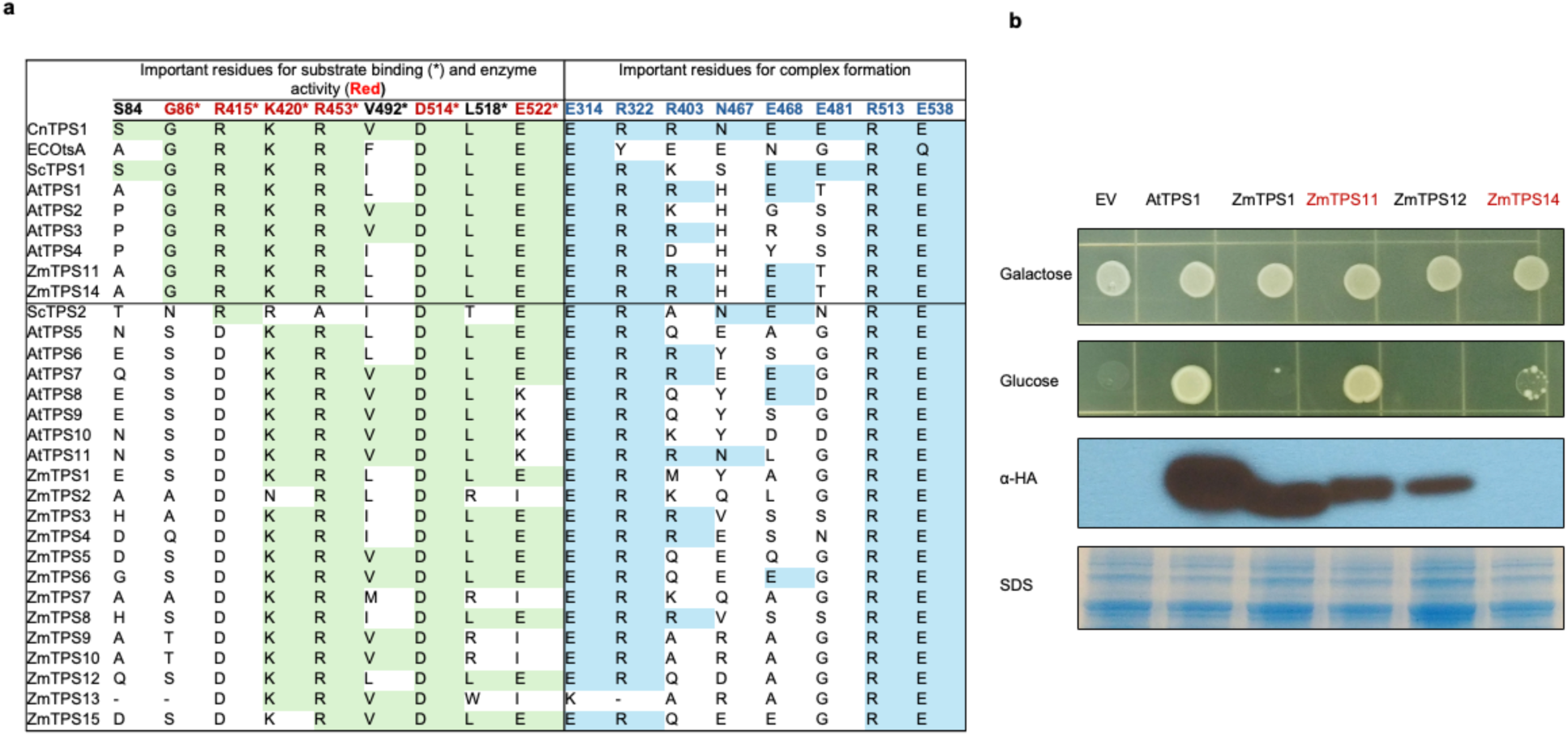
Functional analysis of ZmTPS proteins. (a) Conservation of catalytic and structural residues in TPS homologs across species. Alignment highlights residues critical for substrate binding (green, asterisk), catalytic function (red, asterisk), and complex formation (blue). Alignment is based on *Cryptococcus neoformans* CnTps1 numbering and includes orthologs from *Cryptococcus neoformans* (CnTps1), *E. coli* (EcOtsA), *S. cerevisiae* (ScTPS1-2), *Arabidopsis thaliana* (AtTPS1-11), and *Zea mays* (ZmTPS1-15). Proteins above the horizontal line are considered catalytic, class I TPSs (b) Functional complementation of a *Saccharomyces cerevisiae tps1* mutant. Expression of HA-tagged *AtTPS1*, *ZmTPS11*, or *ZmTPS14* restored growth on glucose, whereas *ZmTPS1* and *ZmTPS12* did not. Western blot against HA-tag confirms protein expression; SDS-PAGE shows equal loading across samples, EV= empty vector.

**Figure S3.**
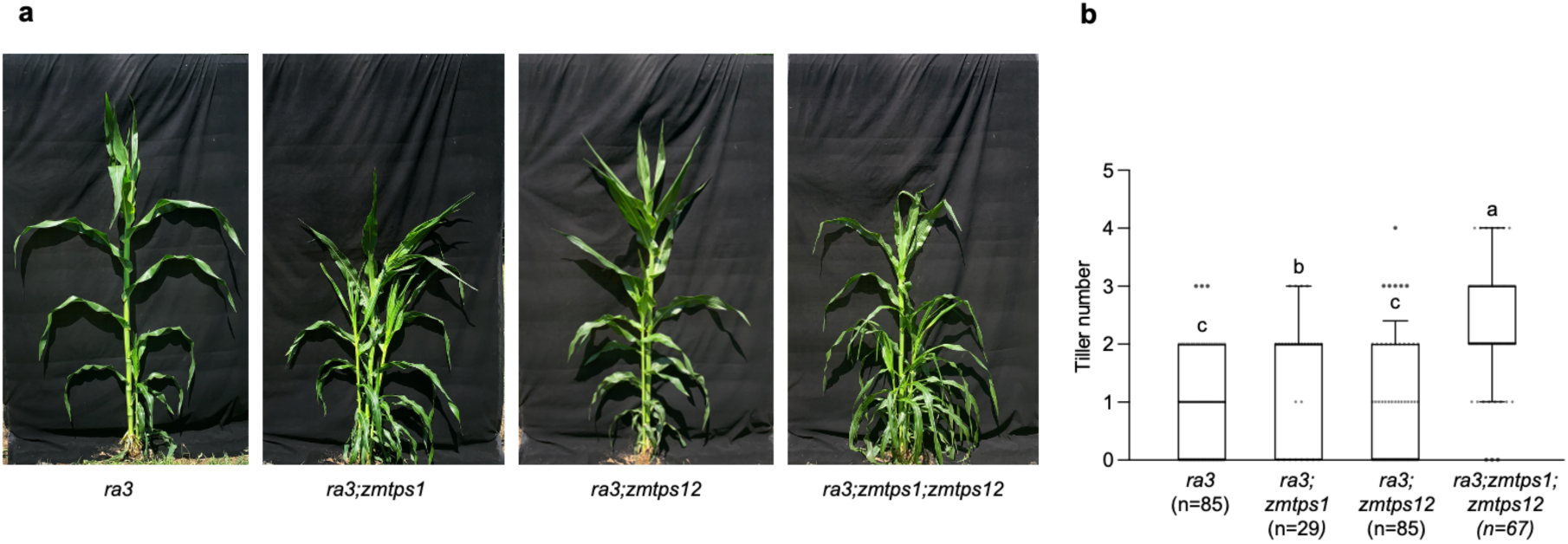
Non-catalytic ZmTPS1 and ZmTPS12 enhance tillering in the *ra3* mutant background. **(a)** Representative plants of the indicated genotypes show increased tillering when *zmtps1* mutations are combined with *ra3*. The *ra3;zmtps1;zmtps12* triple mutant has the most pronounced tiller outgrowth. **(b)** Quantification of tiller number per plant across genotypes. Box plots show median, interquartile range, and outliers; letters indicate statistically significant differences (p < 0.05, ANOVA with post hoc test). Sample sizes are indicated below each genotype. The *ra3;zmtps1;zmtps12* triple mutant shows a significantly higher tiller number compared to all other genotypes.

